# 5-HT Neurons Integrate GABA and Dopamine Inputs to Regulate Meal Initiation

**DOI:** 10.1101/2024.04.26.591360

**Authors:** Kristine M. Conde, HueyZhong Wong, Shuzheng Fang, Yongxiang Li, Meng Yu, Yue Deng, Qingzhuo Liu, Xing Fang, Mengjie Wang, Yuhan Shi, Olivia Z. Ginnard, Yuxue Yang, Longlong Tu, Hesong Liu, Hailan Liu, Na Yin, Jonathan C. Bean, Junying Han, Megan E. Burt, Sanika V. Jossy, Yongjie Yang, Qingchun Tong, Benjamin R. Arenkiel, Chunmei Wang, Yang He, Yong Xu

## Abstract

Obesity is a growing global health epidemic with limited effective therapeutics. Serotonin (5-HT) is one major neurotransmitter which remains an excellent target for new weight-loss therapies, but there remains a gap in knowledge on the mechanisms involved in 5-HT produced in the dorsal Raphe nucleus (DRN) and its involvement in meal initiation. Using a closed-loop optogenetic feeding paradigm, we showed that the 5-HT^DRN^◊arcuate nucleus (ARH) circuit plays an important role in regulating meal initiation. Incorporating electrophysiology and ChannelRhodopsin-2-Assisted Circuit Mapping, we demonstrated that 5-HT^DRN^ neurons receive inhibitory input partially from GABAergic neurons in the DRN, and the 5-HT response to GABAergic inputs can be enhanced by hunger. Additionally, deletion of the GABA_A_ receptor subunit in 5-HT neurons inhibits meal initiation with no effect on the satiation process. Finally, we identified the instrumental role of dopaminergic inputs via dopamine receptor D2 in 5-HT^DRN^ neurons in enhancing the response to GABA-induced feeding. Thus, our results indicate that 5-HT^DRN^ neurons are inhibited by synergistic inhibitory actions of GABA and dopamine, which allows for the initiation of a meal.

## INTRODUCTION

Obesity is currently one of the most significant problems in medicine and is a contributing risk factor for type II diabetes, hypertension, stroke, and coronary heart disease ^1, 2^. The worldwide obesity epidemic continues to expand at an alarming rate while effective treatment developing at a much slower rate. The available therapeutics, like the popular GLP-1 agonists, are promising yet they still have some unwanted side effects, largely due to insufficient understanding of how the brain manages body weight and feeding.

Brain-derived serotonin, also known as 5-hydroxytryptamine (5-HT), is primarily synthesized by neurons in the dorsal Raphe nucleus (DRN) in the midbrain. 5-HT neurons in the DRN (5-HT^DRN^) project to numerous brain regions, including the arcuate of the hypothalamus (ARH) ^3^. The central 5-HT system has become an attractive target for anti-obesity therapies. For example, d-fenfluramine (d-Fen), a pharmacological agent that increases brain 5-HT content ^4^, was among the most effective anti-obesity drugs when it was available, although this therapeutic was withdrawn from the market due to adverse cardiac effects ^5^. In addition, a selective 5-HT 2C receptor agonist (lorcaserin) was approved by the FDA as an anti-obesity medicine in 2012 but was also withdrawn due to an increased cancer risk ^6, 7^. Nevertheless, the 5-HT system remains a promising target for anti-obesity therapeutics; however, the exact mechanism for central 5-HT actions on feeding and body weight regulation remains elusive. A more thorough understanding of the neurocircuitry and molecular pathways associated with 5-HT neurons has the potential to lead to the discovery of superior therapeutic targets for obesity.

Previous studies found that both dopamine (DA) neurons residing in the ventral tegmental area (DA^VTA^ neurons) and GABA neurons in the lateral hypothalamus (GABA^LH^ neurons) are involved in hedonic feeding regulation ^8–10^ and promote feeding ^11–14^. We previously found that DA neurons in the mouse VTA bidirectionally regulate the activity of 5-HT^DRN^ neurons, with weaker stimulation causing DRD2-dependent inhibition and overeating, and stronger stimulation causing DRD1-dependent activation and anorexia ^15^. Further, in a separate study, we found that hunger-driven feeding gradually activates ARH-projecting 5-HT^DRN^ neurons via reducing their responsiveness to inhibitory GABAergic inputs^10^. In the current study, we first used the closed-loop optogenetic approach to activate or inhibit the 5-HT^DRN^→ARH circuit when fasted or fed mice approached food and examined whether this circuit regulates meal initiation. We then combined ChannelRhodopsin-2-Assisted Circuit Mapping (CRACM) and slice electrophysiology to examine how 5-HT^DRN^ neurons respond to GABAergic inputs at fed or fasted conditions. We further used genetic mouse models with the GABA_A_ receptor or DA receptors deleted in 5-HT neurons to reveal how 5-HT^DRN^ neurons integrate GABAergic and DAergic signals to regulate feeding behavior.

## RESULTS

### The 5-HT^DRN^**→**ARH circuit regulates meal initiation

It has been demonstrated that the 5-HT^DRN^→ARH circuit regulates feeding ^10, 16–21^, but it remains unclear whether this circuit regulates meal initiation or termination. Therefore, we used a closed-loop optogenetic approach to determine if the 5-HT^DRN^→ARH circuit regulates meal initiation. To stimulate this circuit, we stereotaxically injected a Cre-dependent AAV vector carrying ChR2 into the DRN of TPH2-CreER mice and implanted an optic fiber for light delivery above the ARH (Fig. 1a and Fig. S1). Mice were fasted for 24 h and then provided with a food pellet in the home cage. Importantly, blue light photostimulation (or yellow light as control) was initiated when mice entered to the “food approach zone” (within 5 cm of the food pellet) and ceased when mice left the zone (Fig. 1b-c). We found that food intake was significantly reduced and the latency to eat was significantly increased with stimulation of the 5-HT^DRN^→ARH circuit upon food approaching, compared to yellow-light control (Fig. 1d and h). There were no significant differences in food bouts, entries to the food approach zone, probability of food intake upon approach, or food bout duration (Fig. 1e-g and i). In a separate cohort of TPH2-CreER mice, we stereotaxically injected a Cre-dependent AAV vector carrying eNpHR3.0 into the DRN and implanted an optic fiber above the ARH to allow for photoinhibition of the 5-HT^DRN^→ARH circuit (Fig. 1a). These mice were tested in a sated state (being fed ad libitum) with the similar closed-loop protocol: yellow light photoinhibition (or blue light as control) was initiated when mice entered to the “food approach zone” and ceased when mice left the zone (Fig. 1b-c). Interestingly, photoinhibition of the 5-HT^DRN^→ARH circuit caused significant increases in food intake, food bouts, probability of food intake upon approach, and bout duration compared to blue light control (Fig. 1j, k, m, and o). We also found a significant decrease in the latency to eat (Fig. 1n) but no significant difference in the entries to the food zone (Fig. 1l). Taken together, these data indicate that the 5-HT^DRN^→ARH circuit plays an important role in meal initiation and food intake.

**Fig. 1:**
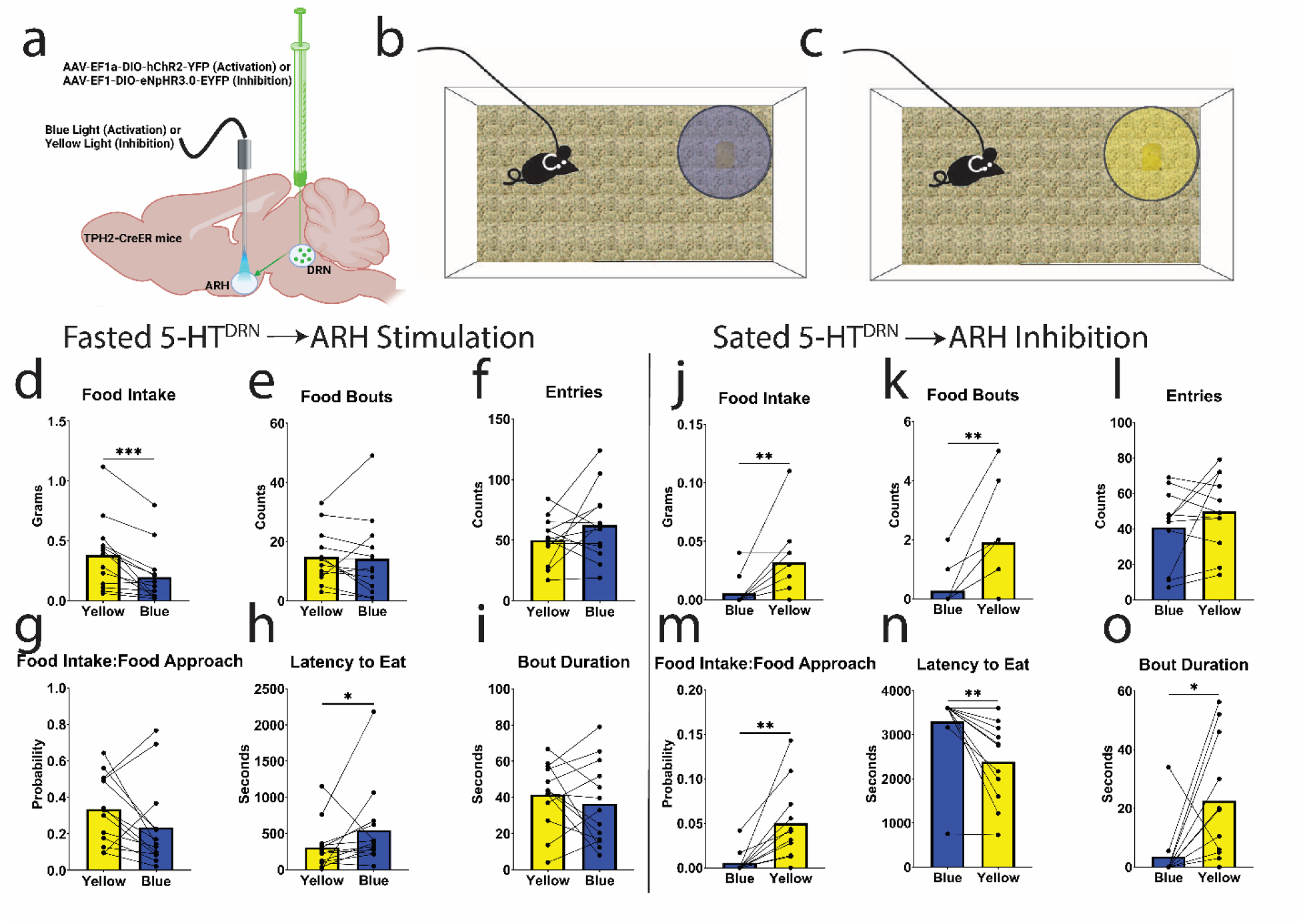
The 5-HT^DRN^→ARH circuit regulates meal initiation. (a) Schematic of viral injection and optic fiber placement. (b) Schematic of blue light stimulation during food approaching. (c) Schematic of yellow light inhibition during food approaching. Results in fasted mice with blue light stimulation of the 5-HT^DRN^→ARH circuit during food approaching (left): (d) food intake, (e) food bouts, (f) entries to the food approach zone, (g) probability of food intake during food approaching, (h) latency to eat, (i) bout duration. Results in sated mice with yellow light inhibition of the 5-HT^DRN^→ARH circuit during food approach (right): (j) food intake, (k) food bouts, (l) entries to the food approach zone, (m) probability of food intake during food approaching, (n) latency to eat, (o) bout duration. Data were analyzed with a paired nonparametric Wilcoxon Test. Data are presented as mean ± SEM, n=11-13 male mice. p < 0.05 *, p < 0.01 **, p < 0.001 ***, p < 0.0001 ****.

**Fig. S1:**
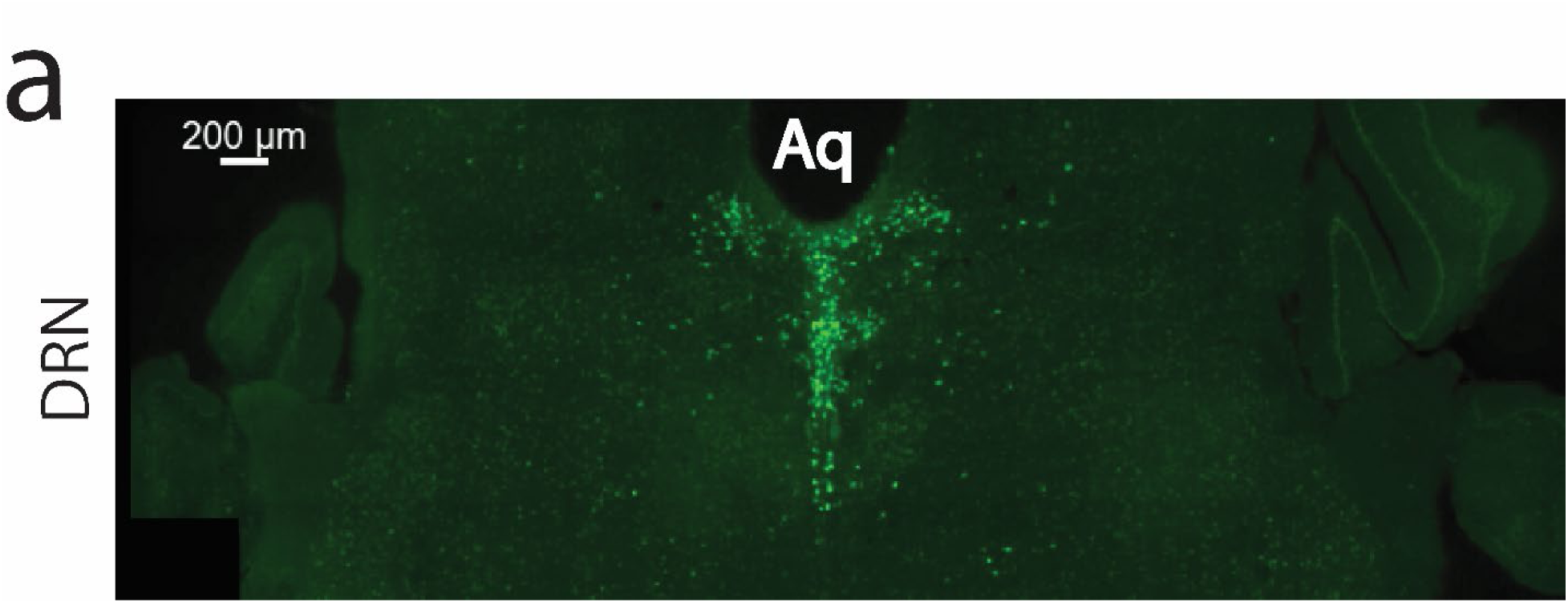
Validation of virus and fiber placement. (a) Representative image of virus placement in the DRN. Scale bar = 200 μm. 3V, third ventricle; Aq, aqueduct; DRN, dorsal Raphe nucleus.

### Responses of 5-HT^DRN^ neurons to GABAergic inputs are regulated by hunger

Since GABAergic neurons within the DRN have local projections and also regulate feeding behavior ^22^, we next examined if 5-HT^DRN^ neurons receive local GABAergic inputs, using the CRACM approach. Briefly, we stereotaxically injected a Flpo-dependent ChR2 into the DRN of TPH2-CreER/Rosa26-LSL-tdTOMATO/Vgat-Flp mice to express ChR2 in Vgat-expressing neurons in the DRN (GABA^DRN^ neurons, Fig. 2a). In these mice, a tamoxifen injection (0.2 mg/g, intraperitoneal) induced Cre recombinase activity in TPH2-expressing (5-HT) neurons and therefore labeled these 5-HT neurons with tdTOMATO. We then prepared brain slices containing the DRN and recorded evoked Inhibitory Post Synaptic Currents (eIPSC) in 5-HT^DRN^ neurons while stimulating GABA^DRN^ neurons with blue light (Fig. 2b). These recordings were performed in the presence of 30 μM DNQX and 30 μM D-AP5 to isolate inhibitory inputs, and we detected time-locked IPSCs evoked with blue light stimulation of GABA^DRN^ neurons (Fig. 2c), indicating that the recorded 5-HT^DRN^ neurons receive inhibitory inputs from local GABA^DRN^ neurons. Then, we eliminated action potential propagation via Na^2+^ and K^+^ channels with the addition of 1 μM TTX and 100 μM of 4-AP, and under this condition, the eIPSCs remained without changes in frequency or amplitude (Fig. 2d and f-g), supporting monosynaptic neurotransmission. Finally, 50 μM of Bicuculline was added which eliminated eIPSCs (Fig. 2e-g), confirming that this input was indeed GABAergic. To further examine how GABAergic inputs to 5-HT^DRN^ neurons are regulated by hunger status, we recorded miniature IPSC (mIPSC) in 5-HT^DRN^ neurons from male and female mice fed ad libitum, fasted for 24 h, or fasted for 24 h followed by refeeding for 2 h (Fig. 2h-l). We found that the amplitude of mIPSCs in 5-HT^DRN^ neurons were significantly increased by 24 h fasting compared to fed mice, which was normalized by the 2-h refeeding (Fig. 2k-l). Notably, no significant differences were observed in mIPSC frequency at these conditions (Fig. S2). Together, these results indicate that 5-HT^DRN^ neurons receive inhibitory inputs partially from the local GABAergic neurons, and their responses to GABAergic inputs can be enhanced by hunger and reduced by feeding. Considering that mIPSC amplitude but not frequency was regulated, we suggest that these regulations are largely mediated by post-synaptic mechanisms.

**Fig. 2:**
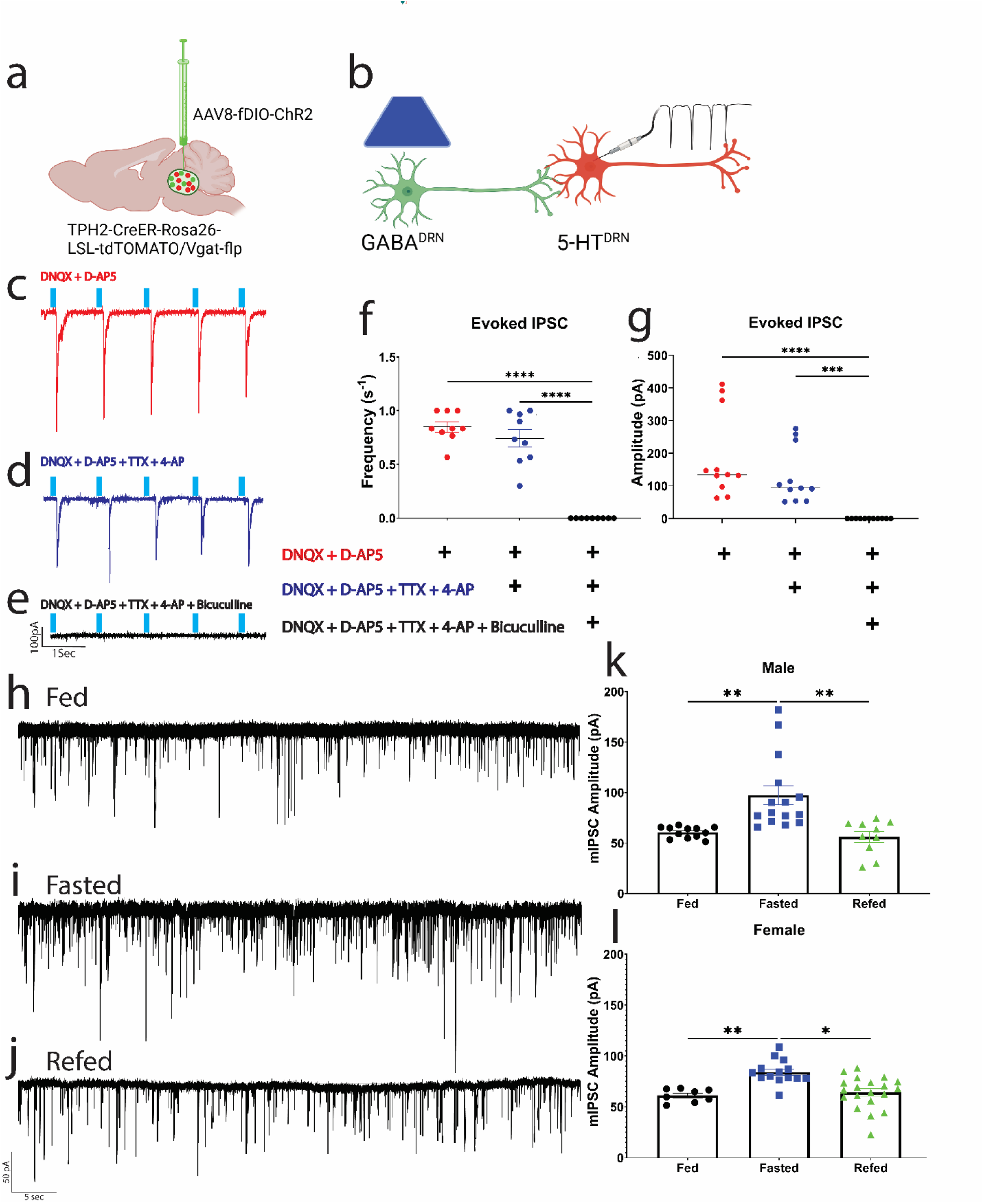
GABA inhibition of 5-HT^DRN^ neurons is regulated by hunger. (a) Schematic representation of viral approach. (b) Schematic of optogenetic electrophysiology recording. (c-e) Representative evoked inhibitory post-synaptic current (eIPSC) traces. (f) Quantification of eIPSC frequency from c-e. (g) Quantification of eIPSC amplitude from c-e. (h-j) Representative miniature inhibitory post-synaptic current (mIPSC) traces. (k-l) mIPSC amplitude quantification of 5-HT^DRN^ neurons from male (k) and female (l) mice in a fed, 24 h fasted and 24 h fasted followed by 2 h refeed (refed) state. Data were analyzed with a two-way ANOVA with post-hoc multiple comparisons. Data are presented as mean ± SEM. For f-g, n=9 cells from 4 male mice. For h-l, n=8-21 cells from 3-5 male and female mice. p < 0.05 *, p < 0.01 **, p < 0.001 ***, p < 0.0001 ****.

**Fig. S2:**
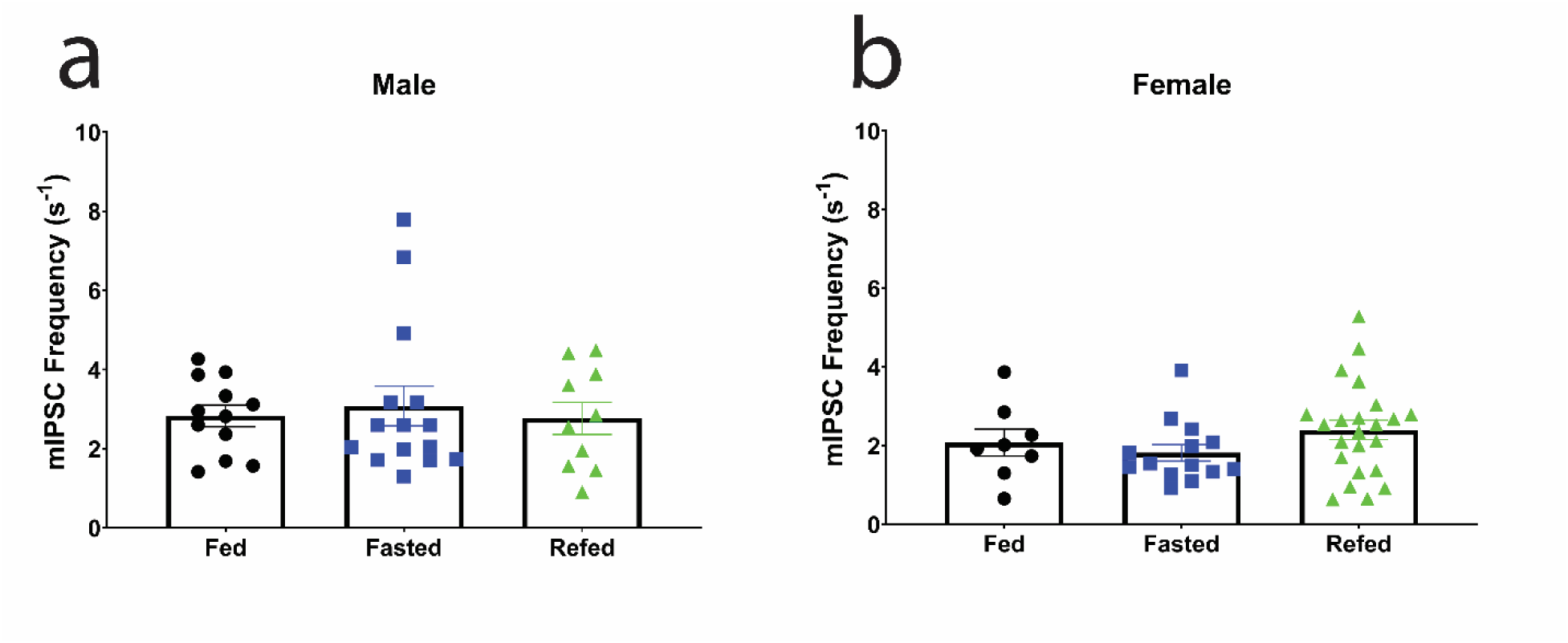
mIPSC frequency in 5-HT^DRN^ neurons. mIPSC frequency in males (a) and females (b) in a fed, 24 h fasted and 24 h fasted followed by 2 h refeed (refed) state. Data were analyzed with a two-way ANOVA with post-hoc multiple comparisons. Data are presented as mean ± SEM, n=8-21 cells from 3-5 male and female mice.

### GABAergic actions in 5-HT^DRN^ neurons regulate meal initiation

It has been reported that GABA signals within the DRN can initiate feeding in sated mice ^22^. Since we observed that GABA^DRN^ neurons provide monosynaptic inhibitory inputs to 5-HT^DRN^ neurons, we next examined whether DRN GABA-induced feeding involves 5-HT signals. To this end, we surgically implanted a cannula above the DRN to allow for direct delivery of a GABA agonist, Muscimol (Fig. 3a). After recovery from the surgery, 0.5 µl of Muscimol was infused at a concentration of 0.5 µg/µl into the DRN of sated mice, and the same volume of sterile saline was used as a control. We found that in control mice, Muscimol significantly increased food intake compared to saline (Fig. 3b and d). We then performed the same DRN Muscimol/saline infusion in mice lacking TPH2 (the rate limiting enzyme for 5-HT synthesis) in the DRN (TPH2^DRN^KO mice). Interestingly, Muscimol had no effect on food intake in TPH2^DRN^KO mice (Fig. 3c and d). Notably, saline-treated TPH2^DRN^KO mice consumed significantly more food compared to saline-treated controls (Fig. 3d), consistent with the notion that 5-HT synthesized by DRN neurons is anorexigenic ^16^. These data indicate the DRN GABA signals within the DRN can trigger food intake and this effect involves 5-HT signals.

**Fig. 3:**
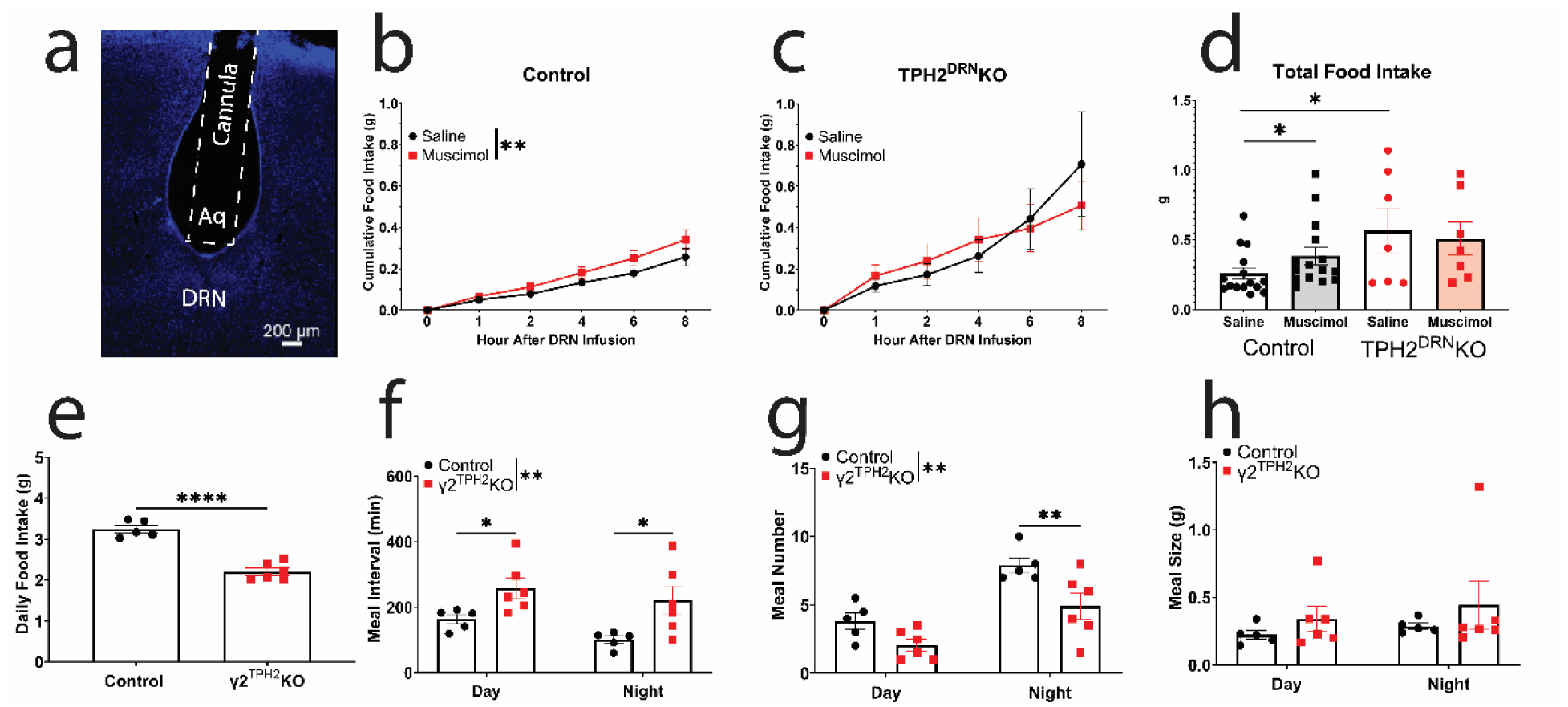
GABAergic actions in 5-HT^DRN^ neurons regulate meal initiation. (a) Representative image of cannula placement above the DRN. (b) Cumulative food intake results in control mice after saline or Muscimol DRN infusion. (c) Cumulative food intake results in TPH2^DRN^KO mice after saline or Muscimol DRN infusion. (d) Cumulative food intake results in control and TPH2^DRN^KO mice after saline or Muscimol DRN infusion. (e) Average daily food intake, (f) meal interval, (g) meal number, and (h) meal size in γ2^TPH2^KO mice and control littermates. Data were analyzed with a two-way ANOVA with post-hoc multiple comparisons, except (d) which was analyzed by two-sided unpaired t-test. Data are presented as mean ± SEM, (b) n=15 male mice, (c) n=8 male mice, (d-g) n=5-6 male mice. p < 0.05 *, p < 0.01 **, p < 0.001 ***, p < 0.0001 ****.

Subsequently, we examined the physiological relevance of GABAergic inputs to 5-HT^DRN^ neurons in feeding regulation. To this end, we generated mice lacking γ2 (a pore-forming subunit of GABA_A_ receptor) in 5-HT neurons (γ2^TPH2^KO mice). We found that γ2^TPH2^KO mice had decreased daily food intake (Fig. 3e). Further detailed meal pattern analysis revealed that loss of γ2 from 5-HT neurons significantly increased meal interval (Fig. 3f), and decreased meal number at night (Fig. 3g). The increased meal interval and decreased meal number suggest that reduced GABAergic signals in 5-HT neurons inhibits meal initiation. Notably, there was no significant change in meal size (Fig. 3h), suggesting that GABAergic signals in 5-HT neurons are not involved in the satiation process. In addition, we found that body weight of γ2^TPH2^KO mice was comparable to control littermates (Fig. S3a), likely due to the observed reduction in energy expenditure in these mice (Fig. S3b-d).

**Fig. S3:**
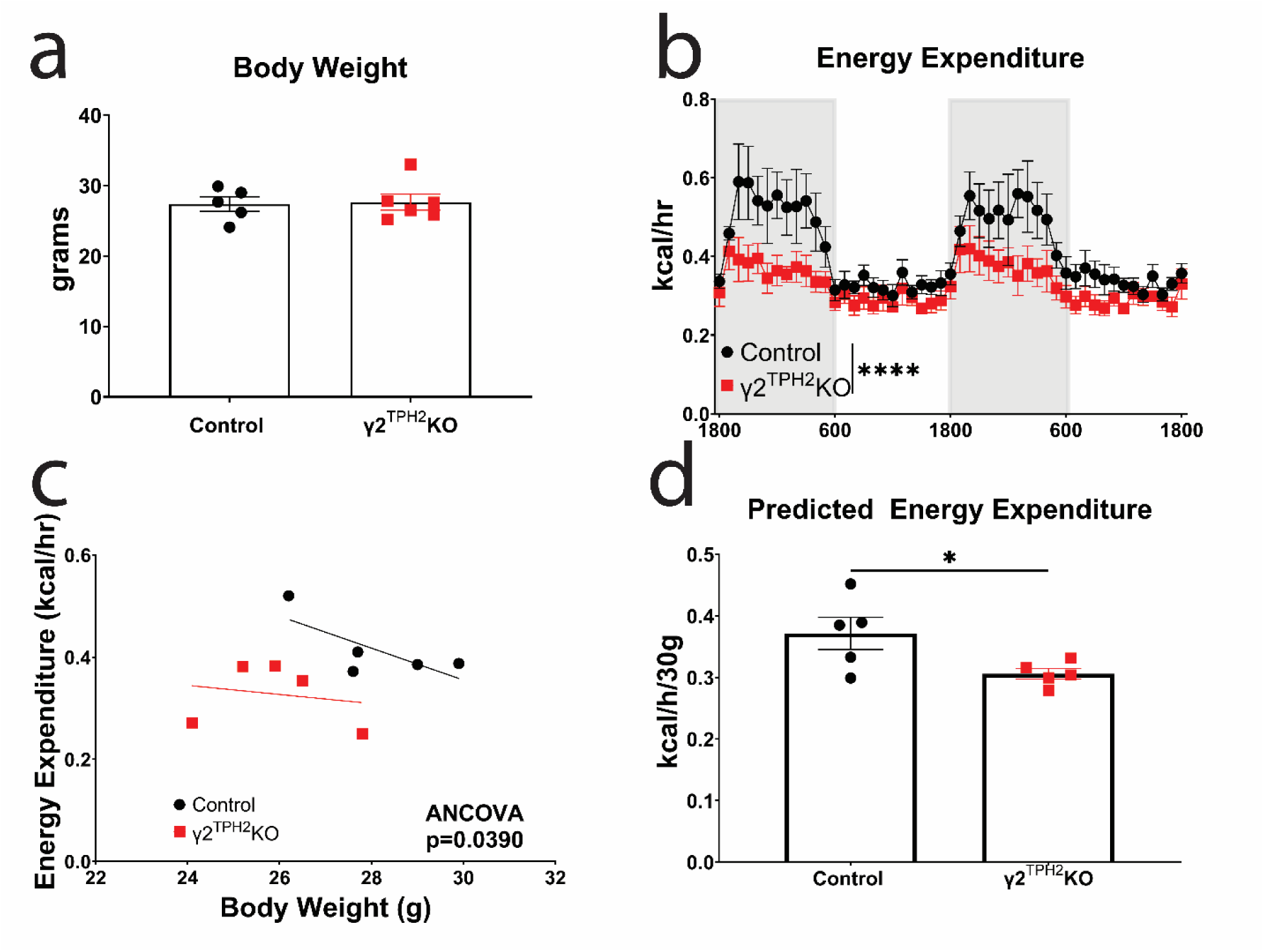
Body weight and energy expenditure in γ2^TPH2^KO. γ2^TPH2^KO and control littermate (a) body weight, (b) hourly energy expenditure, (c) energy expenditure plotted against body weight as a covariate, and (d) predicted energy expenditure for a 30 g mouse. Data were analyzed by (a & d) t-test, (b) two-way ANOVA with post-hoc multiple comparisons, and (c) ANCOVA. Data are presented as mean ± SEM, n=5 male mice. p < 0.05 *, p < 0.01 **, p < 0.001 ***, p < 0.0001 ****.

### DRD2 enhances 5-HT^DRN^ neuronal responses to GABAergic inputs

We have previously reported that 5-HT^DRN^ neurons receive DAergic inputs via DA receptors DRD1 and DRD2 signaling ^15^. Thus, we further explored whether these DA receptors affect how 5-HT^DRN^ neurons respond to GABAergic inputs. Briefly, we performed mIPSC recordings in control, DRD2^TPH2^KO, and DRD1^TPH2^KO male and female mice that were fed ad libitum or fasted for 24 h. In the fed state, the amplitude of mIPSCs in DRD2^TPH2^KO mice (both males and females) was significantly decreased compared to control mice, while the mIPSC amplitude in DRD1^TPH2^KO mice was comparable to that of control mice (Fig. 4a-b). Similarly, we detected significantly reduced mIPSC amplitude in 5-HT^DRN^ neurons from DRD2^TPH2^KO mice after a 24 h fasting, but the mIPSC amplitude in DRD1^TPH2^KO mice was comparable to that of control mice (Fig. 4c-d). In the fed condition, the mIPSC frequency in 5-HT^DRN^ neurons was significantly reduced in DRD1^TPH2^KO female mice, but not in DRD2^TPH2^KO female mice, compared to control mice, whereas in all other conditions tested, the mIPSC frequency remained comparable among the groups (Fig. S4). These data suggest that DRD2 signals within 5-HT^DRN^ neurons are required to enhance their responses to GABAergic inputs, while DRD1 signals have little to no effect. Finally, we examined feeding responses to DRN infusions of Muscimol in DRD2^TPH2^KO mice. While infusions of Muscimol into the DRN significantly increased food intake in control mice, these orexigenic effects were blunted in DRD2^TPH2^KO mice (Fig. 4e). Taken together, these data indicate that DRD2 signals in 5-HT^DRN^ neurons are required for these neurons to fully respond to GABA inhibitory inputs and therefore contribute to the GABA-induced feeding.

**Fig. 4.**
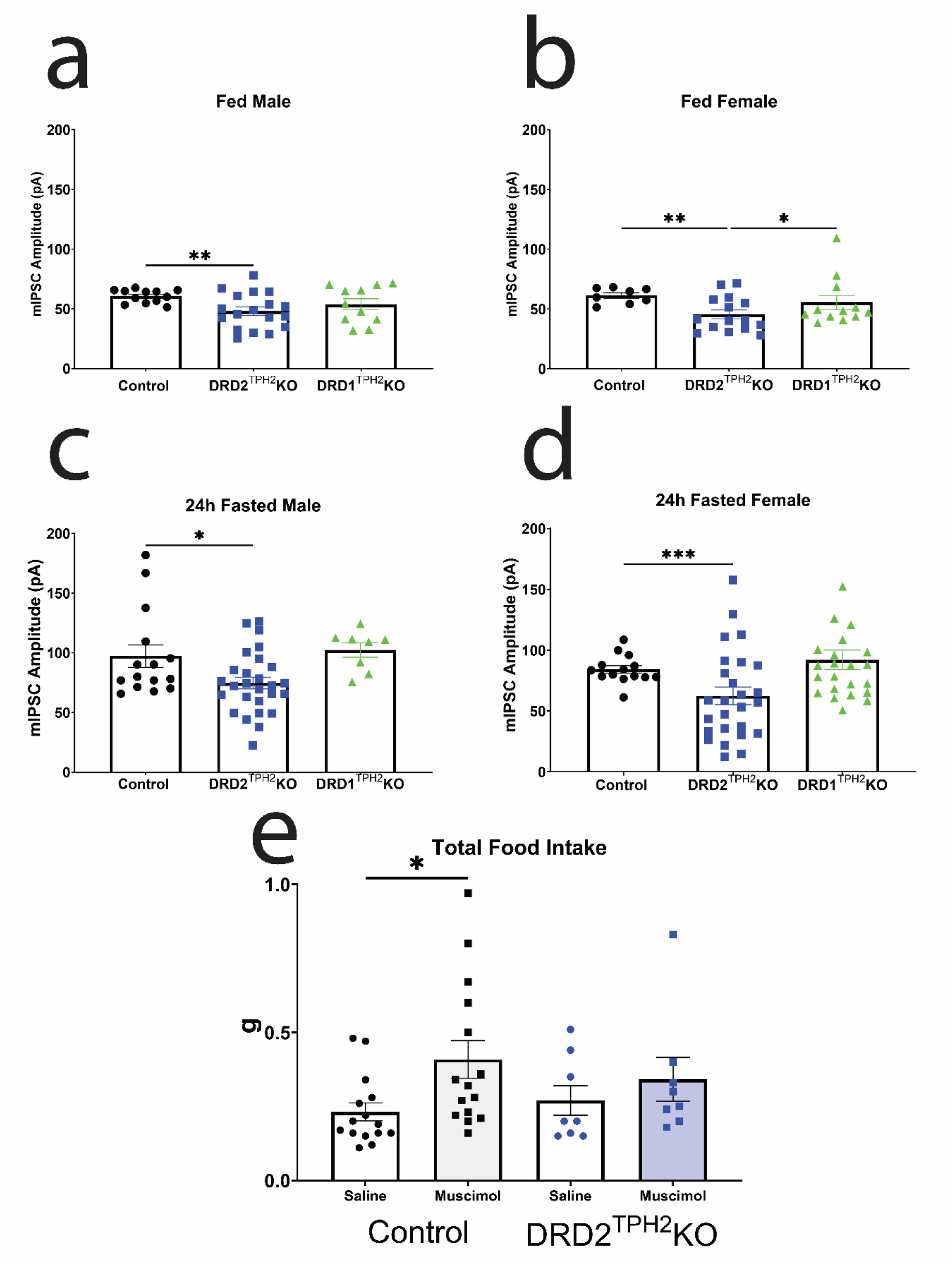
DRD2 enhances 5-HT^DRN^ neuronal responses to GABAergic inputs. mIPSC amplitude in control, DRD2^TPH2^KO, and DRD1^TPH2^KO (a) fed males and (b) fed females. (c) 24 h fasted males and (d) females. (e) Total food intake in control and DRD2^TPH2^KO male mice after saline or Muscimol DRN infusion. Control mIPSC amplitude is the same as presented in Fig. 2k-l. Control Muscimol response is the same as presented in Fig. 3b. Data were analyzed with a two-way ANOVA with post-hoc multiple comparisons. Data are presented as mean ± SEM, (a-d) n=8-21 cells from 3-5 male and female mice, (e) n=8-15 male mice. p < 0.05 *, p < 0.01 **, p < 0.001 ***, p < 0.0001 ****.

**Fig. S4.**
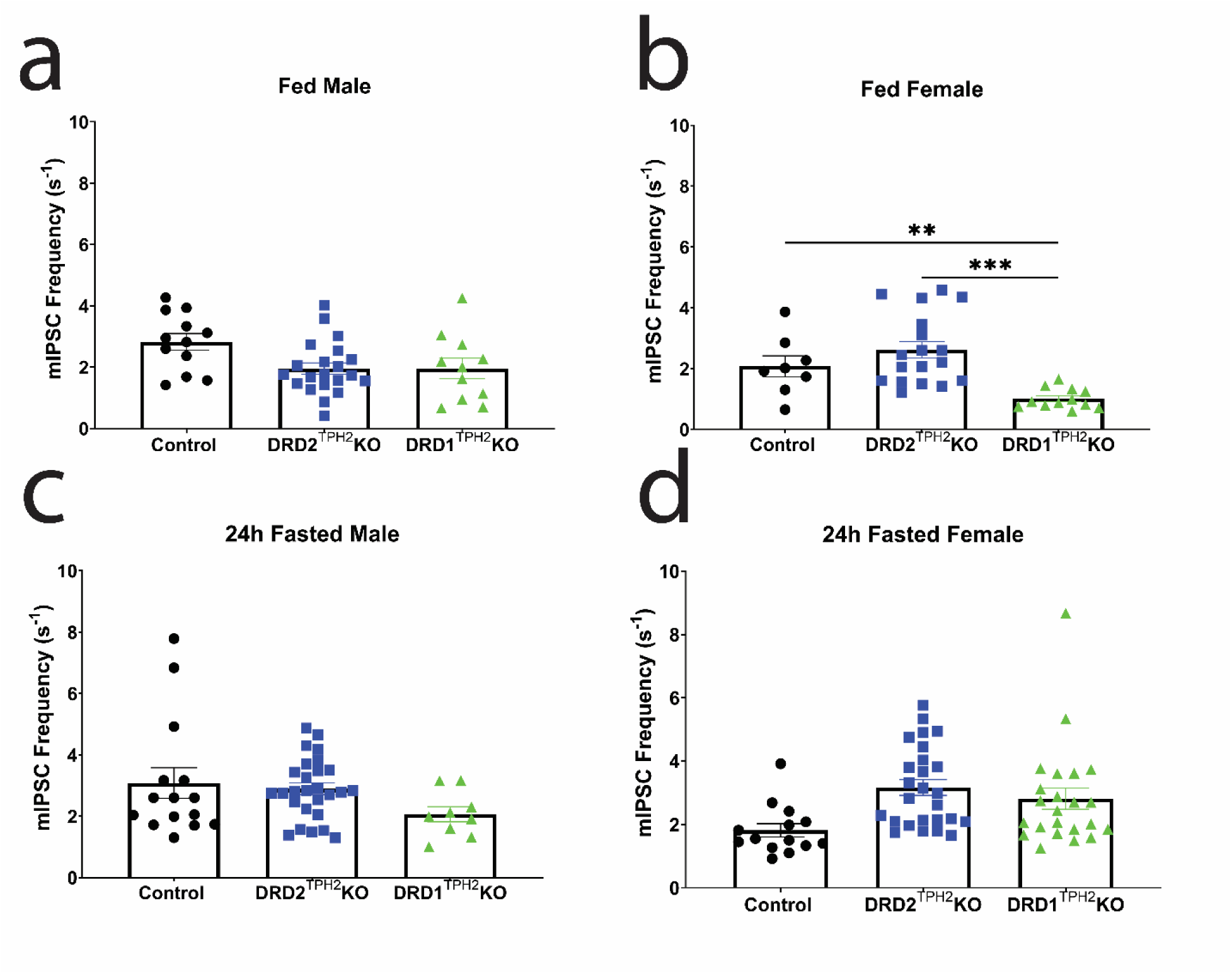
mIPSC frequency. mIPSC frequency in control, DRD2^TPH2^KO, and DRD1^TPH2^KO (a) fed males and (b) fed females. (c) 24 h fasted males and (d) females. Control mIPSC data are the same that are presented in Fig. S2. Data were analyzed with a two-way ANOVA with post-hoc multiple comparisons. Data are presented as mean ± SEM, n= 8-21 cells from 3-5 male and female mice. p < 0.05 *, p < 0.01 **, p < 0.001 ***, p < 0.0001 ****.

## DISCUSSION

Using a closed-loop optogenetic feeding paradigm, we showed that the 5-HT^DRN^→ARH circuit plays an important role in regulating meal initiation and food intake. We further used the CRACM approach coupled with electrophysiology to demonstrate that 5-HT^DRN^ neurons receive inhibitory inputs partially from GABAergic neurons in the DRN, and the 5-HT^DRN^ neuronal response to this GABAergic input can be enhanced by hunger. Further, we found that a direct DRN infusion of the GABA_A_ receptor agonist (Muscimol) can enhance feeding in sated mice, and this effect involves 5-HT signals. Additionally, deletion of the GABA_A_ receptor subunit in 5-HT neurons inhibits meal initiation but has no effect on the satiation process. Finally, we identified the instrumental role of DAergic input via DRD2 in 5-HT^DRN^ neurons in enhancing their response to GABAergic inputs and therefore GABA-induced feeding.

The sources of GABAergic inputs to 5-HT^DRN^ neurons have been established as coming from several regions of the brain including the LH ^23^, VTA ^24^, substantia nigra, amygdala, and preoptic area ^25^. Further, our work corroborates findings which have established that local GABA^DRN^ neurons also contribute to the inhibition of 5-HT^DRN^ neurons ^26, 27^. Inputs from these distinct sources have been found to contribute directly to feeding ^10–12, 28^, including our own work, while others contribute to different behaviors like social defeat ^26^, reward ^27^, and wakefulness ^29^. These GABAergic signals are dynamic during the hunger-satiation transition. For example, in a fed condition, responses to GABAergic signals are decreased, leading to increased 5-HT^DRN^ neuron activity, and subsequently inhibit meal initiation. In a fasted condition, 5-HT^DRN^ neuronal responses to GABAergic signals are increased, leading to inhibition of these 5-HT^DRN^ neurons, which subsequently allow for meal initiation^10^. Importantly, our current work further suggests a post-synaptic (but not pre-synaptic) mechanism for the GABA dynamics in 5-HT neurons, given the changes in mIPSC amplitude but not frequency. In addition to the multiple sources of GABAergic inputs to 5-HT^DRN^ neurons, there are also other neurotransmitters which may mediate the actions of GABA in this context, including glutamate ^22^ as well as DA ^14, 15^.

We have previously shown that 5-HT^DRN^ neurons also receive DAergic inputs, primarily from the VTA ^15^. Interestingly, tonic firing (<10 s^-1^ in frequency) of DA^VTA^ neurons inhibits 5-HT^DRN^ neurons via the DRD2, leading to overeating; on the other hand, phasic bursting (>10 s^-1^ in frequency) of DA^VTA^ neurons activates 5-HT^DRN^ neurons via the DRD1, resulting in anorexia-like behavior in mice ^15, 30^. Notably, the phasic bursting of DA^VTA^ neurons can be triggered by pathological conditions, e.g. the activity-based anorexia, but is rarely seen in mice under normal feeding conditions ^15^. Consistently, we observed that the dynamic changes in GABAergic signals in 5-HT^DRN^ neurons at fed vs. fasted conditions were only affected by the DRD2 deletion while the DRD1 deletion only had a minor effect, presumably because DA^VTA^ neurons predominantly exhibit a tonic firing pattern at this physiological condition. Similar to local GABA inhibition of 5-HT^DRN^ neurons, there are also local DA neurons located in the DRN, which have been reported to regulate social interactions ^31^. Whether these DA neurons within the DRN, or other DA neurons in the brain, can also contribute to the dynamic 5-HT^DRN^ neuronal activity during feeding remains to be explored.

Our results show that the lack of DRD2 in 5-HT neurons reduces mIPSC amplitude, indicating that DRD2 signaling can influence how 5-HT^DRN^ neurons respond to GABAergic inputs. However, the molecular mechanisms for this integration between DAergic and GABAergic signals remain unclear. It was reported that the GABA_A_ receptor can form a complex with DA receptor D5 (DRD5) via the direct binding of the DRD5 carboxy-terminal domain with the second intracellular loop of the GABA_A_ γ2 subunit ^32^. This physical association enables mutually functional interactions between these receptors ^32^. We speculate that the GABA_A_ γ2 receptor subunit and DRD2 may also interact with each other, as one potential mechanism underlying the observed DRD2 influence on GABAergic signaling in 5-HT^DRN^ neurons. Of course, we cannot exclude other possibilities that DRD2 signaling may affect expression of GABA_A_ receptors or their intracellular trafficking ^33^. Additional investigations are warranted to explore these mechanisms.

Further, our data suggest that ARH-projecting 5-HT^DRN^ neurons regulate meal initiation. For example, in fasted mice that naturally engage frequent eating bouts, activation of the 5-HT^DRN^→ARH circuit, only when mice approach the food, can significantly delay the initiation of an eating bout, and reduce total food intake. On the other hand, in sated mice that barely eat, inhibition of this circuit is sufficient to trigger eating with reduced latency and increased bouts. In addition, the loss of GABA_A_ receptor subunit in 5-HT neurons leads to increased meal interval and decreased meal number, without changing the meal size, indicating reduced meal initiation in these mice. The downstream projection from 5-HT^DRN^→ARH neurons involve Agouti-Related Protein (AgRP) neurons in the ARH ^10, 28, 34^. This projection to AgRP^ARH^ neurons is critical as AgRP^ARH^ neurons have been found to be a key regulator of meal initiation and food intake ^28, 34^. In the other direction, the upstream GABAergic input to 5-HT^DRN^ neurons (from the LH ^11^ and locally) plays a clear role in meal initiation, as evidenced by the disrupted meal pattern observed in γ2^TPH2^KO mice.

Limitations in our studies include the unexplained changes in mIPSC frequency in fed female DRD1^TPH2^KO mice but not observed in any other groups. This could be due to differences in estrous cycle at the time of sacrifice. Our data indicate a primarily post-synaptic mechanism due to the consistent change in mIPSC amplitude. However, we could not fully rule out the possibility that a pre-synaptic mechanism could also contribute to this proposed circuit. An additional limitation of our study is that some studies (electrophysiology) were repeated in females with similar outcome as males, but not all studies are performed in both sexes. Additional experiments including females needs to be done, taking the estrous cycle into account, to determine if there are any sex differences involved in this circuit.

In summary, our data support a model that in a state of hunger, 5-HT neurons in the DRN are inhibited by synergistic actions of GABA and DA (via the DRD2), which allows the initiation of a meal. As animals feed and satiety is reached, the inhibitory signals in 5-HT^DRN^ neurons are reduced, leading to increased 5-HT release to inhibit feeding via projections to the ARH (Fig. 5).

**Fig. 5.**
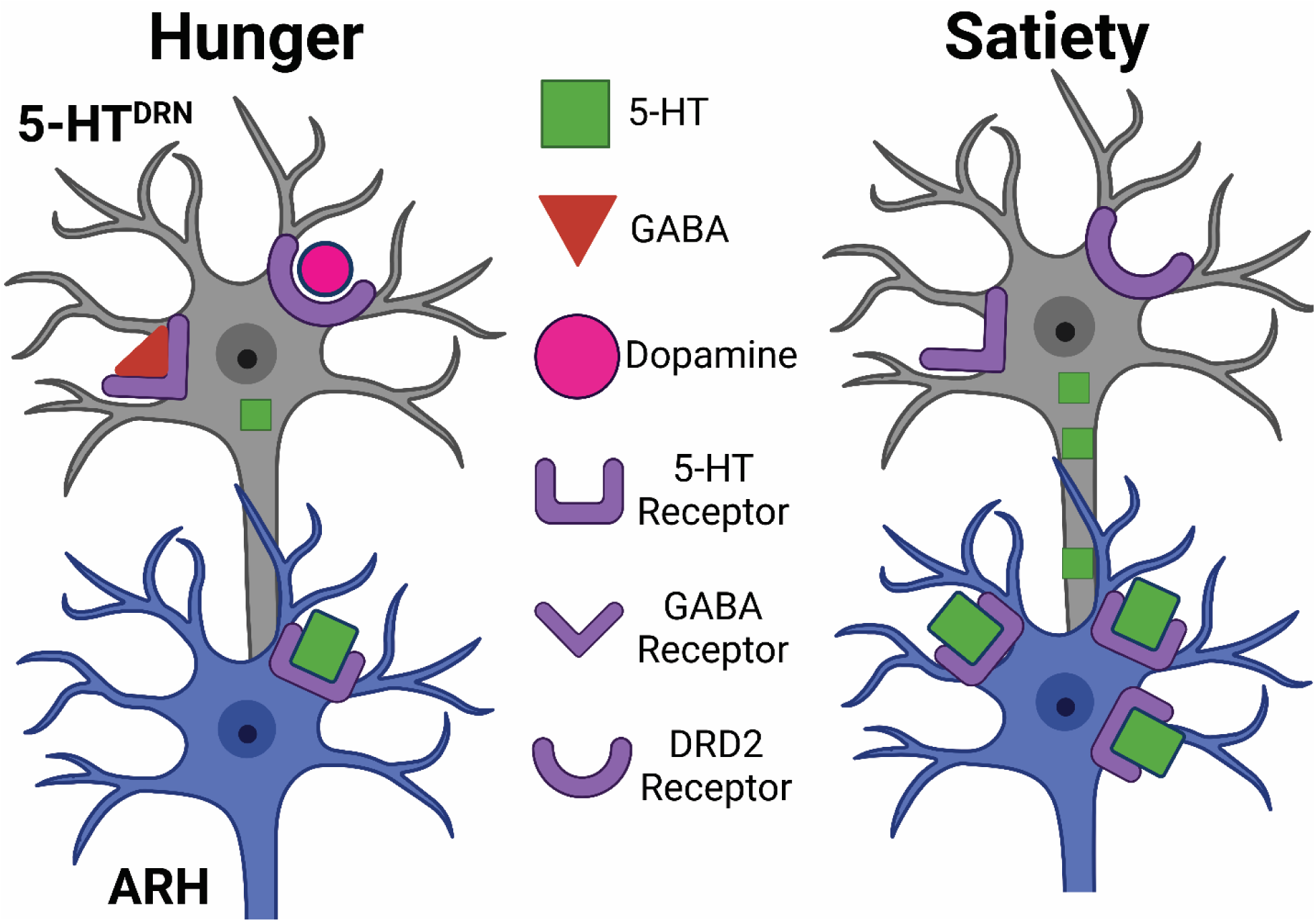
Hypothetic model. The proposed hypothesis of this paper indicates that in a state of hunger, 5-HT neurons in the DRN which project to the ARH, are simultaneously inhibited by both GABA and DA (via the DRD2 receptor) to initiate a meal. Once satiety is reached, 5-HT neurons in the DRN are no longer inhibited and therefore more 5-HT is released into the ARH, reducing food intake.

## METHODS

### Mice

Care of all animals and procedures were approved by Baylor College of Medicine Institutional Animal Care and Use Committees. Mice were housed in a temperature-controlled environment at 22–24 °C, using a 12-h light, 12-h dark cycle. The mice were fed with regular chow (Pico Lab, LabDiet, 5V5R). Water was provided ad libitum.

Multiple lines of transgenic mice were used in the current study, as summarized in **Table 1**. TPH2-CreER mice were purchased from Jackson Laboratory (016584) that express tamoxifen-inducible Cre recombinase selectively in 5-HT neurons, as we validated previously ^15, 35^. Furthermore, we crossed TPH2-CreER allele onto γ2^fl/fl^ mice ^36^ (Jackson Laboratory, #019083) a model we previously validated ^10^. In addition, we crossed Drd1^fl/fl^ mice (Jackson Laboratory, 025700) ^37^ and TPH2-CreER mice to generate Drd1^fl/fl^/TPH2-CreER mice (DRD1^TPH2^KO) and their littermate controls (Drd1^fl/fl^). Similarly, we crossed Drd2^fl/fl^ mice (Jackson Laboratory, 020631) ^38^ and TPH2-CreER mice to generate Drd2^fl/fl^/TPH2-CreER mice (DRD2^TPH2^KO) and their littermate controls (Drd2^fl/fl^). DRD1^TPH2^KO and DRD2^TPH2^KO have been used and validated previously by our lab ^15^. All mice received tamoxifen injections (0.2 mg/g, intraperitoneal) at 8 weeks of age to induce Cre activity. All the breeders have been backcrossed to C57Bl6j background for >12 generations.

**Table 1.**
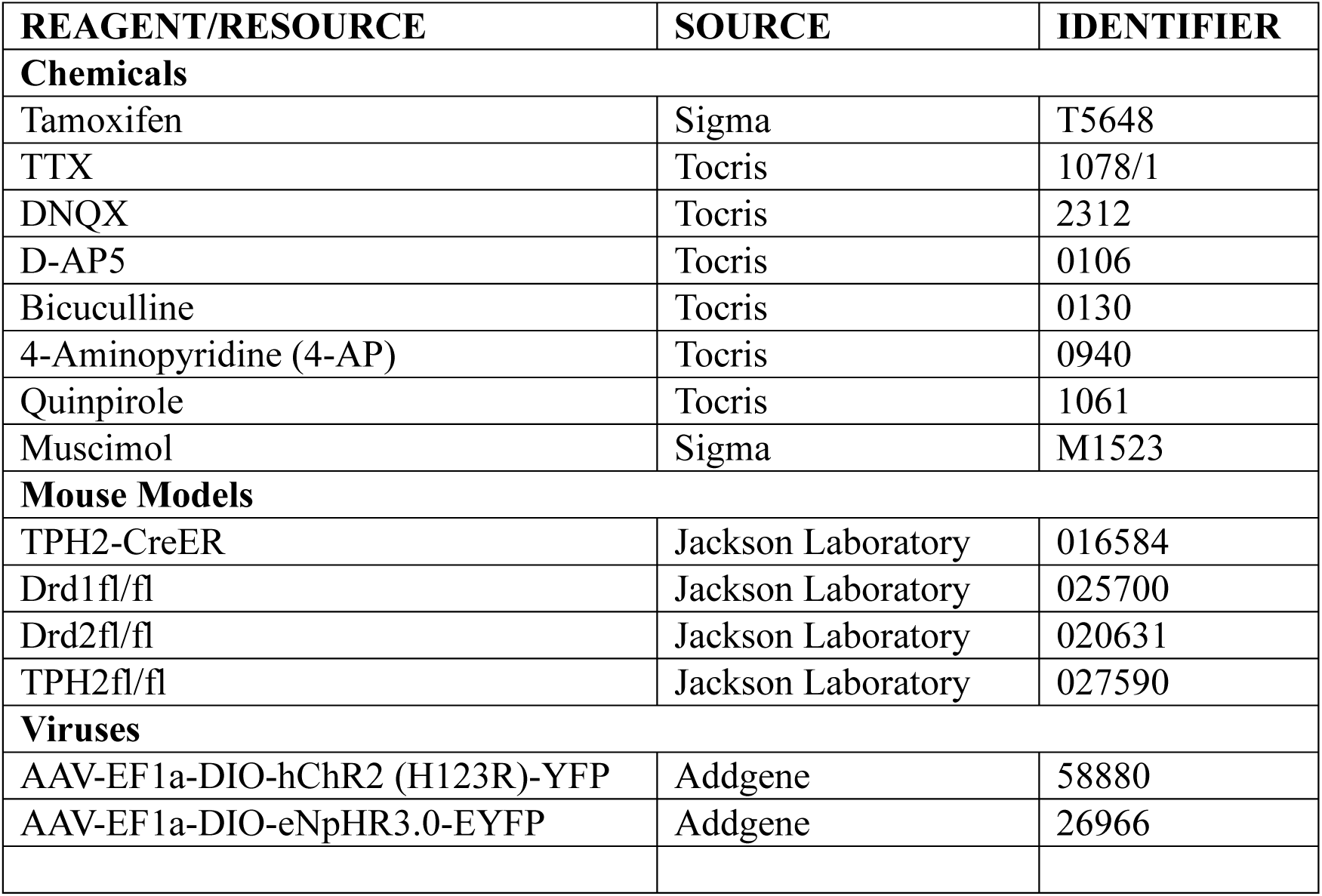
List of Resources.

| REAGENT/RESOURCE | SOURCE | IDENTIFIER |
| --- | --- | --- |
| **Chemicals** |  |  |
| Tamoxifen | Sigma | T5648 |
| TTX | Tocris | 1078/1 |
| DNQX | Tocris | 2312 |
| D-AP5 | Tocris | 0106 |
| Bicuculline | Tocris | 0130 |
| 4-Aminopyridine (4-AP) | Tocris | 0940 |
| Quinpirole | Tocris | 1061 |
| Muscimol | Sigma | M1523 |
| **Mouse Models** |  |  |
| TPH2-CreER | Jackson Laboratory | 016584 |
| Drd1<sup>fl/fl</sup> | Jackson Laboratory | 025700 |
| Drd2<sup>fl/fl</sup> | Jackson Laboratory | 020631 |
| TPH2<sup>fl/fl</sup> | Jackson Laboratory | 027590 |
| **Viruses** |  |  |
| AAV-EF1a-DIO-hChR2 (H123R)-YFP | Addgene | 58880 |
| AAV-EF1a-DIO-eNpHR3.0-EYFP | Addgene | 26966 |

For studies including female mice, estrous phase was recorded at time of sacrifice. Data were analyzed between females by estrous phases; since no significant differences were observed among different estrous phases, female data were pooled regardless of the estrous phases.

### Closed-Loop Optogenetics

Two weeks after tamoxifen induction, mice were stereotaxically injected with either AAV-EF1a-DIO-hChR2 (H123R)-YFP (4×1012 GC per ml) or AAV-EF1-DIO-eNpHR3.0-EYFP (4×1012 GC per ml) into the DRN with coordinates AP: −4.65, ML: 0, DV: −3.4 from the bregma (250 nl) and AP: −4.65, ML: 0, DV:-3.1 from the bregma (250 nl). In the same surgery, an optic fiber (RWD Fiber Optic Cannula with 1.25mm Ceramic Ferrule, 200um core, 0.39NA, 6mm long) was implanted above the ARH with coordinates AP: −1.7, ML: -.30 DV: −5.9 from the bregma. After 4 weeks of recovery, mice were acclimated to optogenetics fibers, testing room, and glass petri dish four times for 10 minutes each prior to experimentation. For stimulation studies, mice were fasted for 24 h prior to testing; for inhibition studies, mice were tested in a fed state. All mice were moved to a new home cage on the morning of the experiment. Mice were allowed to acclimate to testing room for a minimum of 1 h prior to testing. All mice were tested 1 week apart, and the order of light exposure (blue and yellow light) and testing order was randomized each week.

Mouse bedding was removed, and a glass petri dish (10 cm) was placed in the corner of the home cage. A single piece of weighed chow was placed in the center of the petri dish and bedding was placed inside the dish, surrounding the chow. The optogenetic fiber was coupled to the implanted fiber and mice were allowed to feely roam their home cage for 1 h. Optogenetic blue light (473 nm, 10 ms per pulse, 20 s^-1^ frequency) or yellow light (589 nm, 10 ms per pulse, 20 s^-1^ frequency) was provided once both front paws entered the food zone (inside the petri dish) and remained on until the front paws exited the food zone. The chow was weighed again at the end of the hour. Fasted mice were immediately provided access to food upon the end of the experiment.

Videos of the mouse behavior was manually analyzed by three investigators. Analysis included food intake, calculated by the weight of food pellet at start minus the weight of food pellet at the end of the experiment. Food bout time was defined as a period of continuous eating lasting longer than one bite or one second. Number of entries to the food approach zone was recorded. The probability of food intake to food approach was calculated by food bouts divided by entries to the food approach zone. The latency to first bite was defined as the time from the start of the experiment for the mouse to take first bite of food. The bout duration was calculated as the average period spent in an eating bout.

All mice were perfused to validate the accuracy of the viral injection as well as optic fiber placement. Mice with inaccurate viral or fiber placement were excluded from the dataset. The order of blue and yellow light stimulation was randomized to avoid potential sequence effects.

### Electrophysiology

Mice were sacrificed when fed ad libitum, after a 24 h fasting, or after a 24 h fasting followed by a 2 h refeeding. The estrous cycle was checked in female mice prior to sacrifice. Mice were deeply anesthetized with isoflurane and transcranial perfused with a modified ice-cold sucrose-based cutting solution (pH 7.3) containing 10 mM NaCl, 25 mM NaHCO3, 195 mM sucrose, 5 mM glucose, 2.5 mM KCl, 1.25 mM NaH2PO4, 2 mM Na-pyruvate, 0.5 Mm CaCl2, and 7 mM MgCl2, bubbled continuously with 95% O2 and 5% CO2. The mice were then decapitated, and the entire brain was removed and immediately submerged in the ice-cold cutting solution. Coronal brain slices (220 mm) containing the DRN were cut with a Microm HM 650 V vibratome (Thermo Scientific) in oxygenated cutting solution. Slices were then incubated in oxygenated artificial CSF (aCSF; 126 mM NaCl, 2.5 mM KCl, 2.4 mM CaCl2, 1.2 mM NaH2PO4, 1.2 mM MgCl2, 11.1 mM glucose, and 21.4 mM NaHCO3, balanced with 95% O2/5% CO2, pH7.4) to recover ∼30 min at 32°C and subsequently for 1hr at room temperature before recording. Slices were transferred to a recording chamber and allowed to equilibrate for at least 10 min before recording. The slices were superfused at 34°C in oxygenated aCSF at a flow rate of 1.8-2 ml/min. tdTOMATO-labeled TPH2 neurons were visualized using epifluorescence and IR-DIC imaging on an upright microscope (Eclipse FN-1, Nikon) equipped with a movable stage (MP-285, Sutter Instrument). Patch pipettes with resistances of 3-5 MΩ were filled with intracellular solution (pH 7.3) containing 140 mM CsCl, 10 mM HEPES, 5 mM MgCl2, 1 mM BAPTA, 5 mM (Mg)ATP, and 0.3 mM (Na)2GTP (pH 7.30 adjusted with NaOH; 295 mOsm/kg). Recordings were made using a MultiClamp 700B amplifier (Axon Instrument), sampled using Digidata 1440A and analyzed offline with pClamp 10.3 software (Axon Instruments). Series resistance was monitored during the recording, and the values were generally <10 MΩ and were not compensated. The liquid junction potential was +12.5 mV and was corrected after the experiment. Data were excluded if the series resistance increased dramatically during the experiment. Currents were amplified, filtered at 1000 s^-1^, and digitized at 20000 s^-1^. Voltage clamp was utilized to record current with a holding potential of −70 mV in the presence of 1μM TTX, 30μM D-AP5 and 30μM DNQX to isolate inhibitory currents. mIPSC frequency and amplitude were analyzed in 60 second bins during a period of stable recording. For CRACM studies IPSCs were evoked by blue light (473 nm, 10 ms per pulse, 1 s^-1^ frequency). 30μM DNQX and 30μM D-AP5 were first perfused to isolate inhibitory input. 1μM TTX and 100μM of 4-AP were added to eliminate action potential propagation via Na^2+^ and K^+^ channels. Finally, 50μM of Bicuculline was added to eliminate GABAergic signaling.

### Muscimol Studies

Eight-week-old mice were stereotaxically implanted with a guide cannula above the DRN with coordinates: AP: 4.65, ML: 0, DV:-3.0 from the bregma. The opening of the cannula projected 2 mm ventral to the surface of the skull. All mice were allowed to recover for 2 weeks before DRN infusion of either sterile saline or the GABA_A_ receptor agonist (Muscimol). Muscimol and saline infusions were done in the same mice in different trials. Muscimol was made according to manufacturer’s instructions and 0.5 µl total volume was infused at concentration of 0.5 mg/ml based on previous publications ^22^. Infusions were randomized and separated by a minimum of one week. All mice were perfused to validate the accuracy of the cannula placement. Mice with inaccurate placement were excluded from the dataset.

### Metabolic Phenotyping

Body weight and food intake were collected weekly starting two weeks after tamoxifen injection. Mice and their controls were singly housed 1 week before tamoxifen injection, at 8 weeks of age; mice were fed ad libitum with a regular chow diet (5V5R-Advanced Protocol PicoLab Select Rodent 50 IF/6F) from weaning to 18 weeks of age. Energy expenditure measurements were performed in temperature-controlled (23 °C) cabinets containing 16 TSE PhenoMaster metabolic cages, to which mice were acclimated for a minimum of 2 days. Data collected from days 3–5 were used for analyses and energy expenditure was analyzed using the online CalR tool (A Web-based Analysis Tool for Indirect Calorimetry Experiments ^39^).

### Statistics

All data were tested for normality using a Shapiro-Wilk test and outliers were identified using a Grubb’s test. If data was normally distributed, comparisons were made with a parametric t-test. If data were not normally distributed, comparisons were made with a non-parametric t-test. For comparisons among groups, a Two-way ANOVA was used with post-hoc multiple comparisons. The minimal sample size was predetermined by the nature of experiments. The data are presented as mean ± SEM and/or individual data points. Statistical analyses were performed using GraphPad Prism to evaluate normal distribution and variations within and among groups. Methods of statistical analyses were chosen based on the design of each experiment and are indicated in figure legends. p < 0.05 was considered statistically significant.

## ACKNOWLEDGEMENTS

The investigators were supported by grants from the USDA/CRIS (51000-064-01S to YX, 3092-51000-062-04(B)S to CW), American Heart Association (23POST1030352 to HL), NIH (F32DK134121 to KMC; R01DK120858 to YX, R01DK131446 to QT and OZG).

## AUTHOR CONTRIBUTIONS

KC and YX conceived the project, experimental design and wiritng the manuscript. KC performed the procedures, data acquisition and analyses. HW and FS assisted in the optogenetics video analysis and muscimol data aqusition. HW also assisted with virus validation. YL, MY, YD, QL, XF, MW, YS, OZG, YY, LT, Hesong Liu, Hailan L, NY, JCB, JH, MEB, SVJ, and YY contributed to the generation of study mice and data discussion. QT, BRA, CW and YH were involved in study design and data discussion.

